# Deep Learning-Driven Fragment Ion Selection for Improved Quantification in MS based Proteomics

**DOI:** 10.64898/2026.01.08.698341

**Authors:** Duc Tung Vu, Georg Wallmann, Marvin Thielert, Enes Ugur, Marc Oeller, Maximilian Zwiebel, Constantin Ammar, Matthias Mann

**Affiliations:** Department of Proteomics and Signal Transduction Max Planck Institute of Biochemistry, Martinsried Germany

**Keywords:** Proteomics, bioinformatics, quantification, open source

## Abstract

Quantitative proteomics relies on accurate selection of fragment ions for quantification, yet most current algorithms apply simple strategies such as median intensity or single quality filters. Modern data-independent acquisition (DIA) searches generate rich features such as fragment ion correlations, retention time and many others that could be leveraged to assess fragment quality. We introduce QuantSelect, a novel strategy to select optimal fragments by systematically integrating these features via self-supervised deep learning. QuantSelect uses a regularized, weighted-variance loss on intensity traces normalized via our directLFQ algorithm. This allows learning a fragment quality score without ground truth labels, enabling on-the-fly training on label-free DIA datasets. Integrated within our alphaDIA pipeline, QuantSelect significantly improves quantitative accuracy and in some cases substantially corrects protein intensity estimation. Sensitivity in differential expression improved by 68% in a mixed-species benchmarking dataset and by 18% in single-cell data. QuantSelect provides a practical framework for data-driven fragment selection that improves accuracy, precision and downstream inference in DIA proteomics.

## Introduction

Quantitative proteomics has emerged as an indispensable approach for systems-level analysis of protein expression, enabling comprehensive characterization of cellular states and their perturbations. The analytical capabilities of modern mass spectrometry instruments allow simultaneous measurement of thousands of proteins and their abundance changes across diverse biological conditions. These quantitative measurements have proven invaluable across clinical research, single-cell proteomics, and spatial proteomics applications (1). Recent technological advancements in data-independent acquisition (DIA) proteomics have dramatically expanded experimental throughput, now enabling the analysis of hundreds of samples per day while maintaining deep proteome coverage (2, 3).

However, DIA experiments generate increasingly complex data characterized by substantial noise and signal interference from co-eluting and co-fragmenting peptide species (4), posing a critical challenge in quantitative proteomics. Accurate quantification depends on reliable fragment-ion selection, yet interference and variable fragment quality often limit precision and accuracy. Addressing these limitations is essential to realize the full potential of high-throughput DIA proteomics in biological discovery and clinical research.

Multiple proteomics domains have been revolutionized by deep learning (5), including peptide spectrum match (PSM) rescoring through ensemble neural networks in DIA-NN (6), prediction of spectral intensities, retention times and ion mobilities (7–11), and de novo sequencing (12–15). However, these advances have not yet been applied to quantitative analysis due to a critical challenge: the absence of reliable ground truth data for protein abundances, which prevents conventional supervised learning approaches. While DIA generates numerous features that could inform quantitative quality assessment (16), how to optimally select the promising and reject the low-quality features remains unresolved.

Here, we present QuantSelect, a deep learning framework that focuses on integrating combinations of quality features, while preserving unmodified intensity values for label-free quantification in a self-supervised manner. QuantSelect solves the missing ground truth problem by leveraging the principle that median intensities of high-quality proteins approximate true abundance values in the majority of cases. It achieves this via a regularized weighted variance loss function. This approach enables effective online training directly on every new dataset, providing a flexible framework that allows users to simply add diverse fragment characteristics such as XIC correlation, ΔRT, m/z error, or any other combination of scores that could indicate fragment quality. QuantSelect then identifies optimal scoring feature combinations. Our approach builds upon our earlier work on directLFQ (17) and in particular its efficient intensity trace normalization to enable variance estimation and derive accurate protein quantities.

Modern search engines including alphaDIA (18), DIA-NN (6), and MaxDIA (19) employ different filtering strategies to exclude low-quality fragments that could compromise quantitative accuracy. For instance, alphaDIA implements extracted ion chromatogram (XIC) correlation filter to identify and remove fragments with poor quantitative properties. Further work has explored fragment selection by careful modeling of the underlying raw data (20), developing novel statistical scores (21) or machine learning (22). All these methods provide novel features that can be predictive of quantitative quality, however, the question on how to optimally integrate different features remains open. With QuantSelect, we introduce a deep learning-based approach for integrating quality features into a unified quality score, providing an intelligent preprocessing step for fragment ion selection, applied before LFQ calculation, enhancing both quantification accuracy and the reliability of downstream data analysis.

To benchmark QuantSelect, we utilized alphaDIA, our recently developed open-source proteomics search engine that achieves performance parity with established frameworks (6, 19, 23). Its modular architecture and comprehensive data output made seamless integration of QuantSelect possible, enabling controlled comparison between ion selection schemes while maintaining all other parameters constant. This is not true for most other frameworks and we therefore focused our comparative analysis on alphaDIA’s implementation versus QuantSelect-enhanced add-on processing.

For validation and benchmarking of QuantSelect, we chose a mixed-species dataset consisting of *H. sapiens*, *S. cerevisiae* and *E. coli* peptides spiked at three defined ratios (high, medium, and low fold change (FC)) (24), acquired on the state-of-the-art Thermo Scientific Orbitrap Astral mass spectrometer (25, 26). We first compared QuantSelect against a simple variance-based filtering approach, ensuring accurate representation of quality features for both high and low-quality fragments. Subsequently, we evaluated QuantSelect’s performance against alphaDIA’s native XIC_corr filtering approach, assessing quantitative accuracy, precision, and sensitivity in differential expression analysis.

QuantSelect is directly integrated into the alphaDIA search engine. It is an open-source Python package that is freely available at https://github.com/MannLabs/quantselect and can be easily applied to arbitrary data formats.

### Experimental Procedures

### Experimental Design

#### Single-cell sample preparation

Naive E14 mouse embryonic stem cells (mESCs) were cultured in serum-free medium consisting of N2B27 [50% Neurobasal medium (Life Technologies), 50% DMEM/F12 (Life Technologies)] supplemented with 2i [1 μM PD0325901 and 3 μM CHIR99021 (Axon Medchem, Netherlands)], 1000 U/ml recombinant leukemia inhibitory factor (LIF; Millipore), 0.3% BSA (Gibco), 2 mM L-glutamine (Life Technologies), 0.1 mM β-mercaptoethanol (Life Technologies), N2 supplement (Life Technologies), B27 serum-free supplement (Life Technologies), and penicillin-streptomycin (100 U/ml and 100 μg/ml, respectively; Sigma).

Formative EpiLCs were derived by differentiating naive mESCs for 48 h in the same serum-free medium without 2i, LIF, and BSA, but supplemented with 10 ng/ml Fgf2 (R&D Systems), 20 ng/ml Activin A (R&D Systems), and 0.1x Knockout Serum Replacement (KSR; Life Technologies) (differentiation medium). Primed EpiSCs were derived by differentiating naive mESCs for 7 days in differentiation medium. All pluripotent stem cell models were cultured on 0.2% gelatin-treated 6-well plates. Medium was changed once after 24 h for EpiLCs (and after 24 h and 3 days for EpiSCs), and cells were harvested after 48 h and 7 days, respectively. Cells were tested negative for Mycoplasma contamination by PCR. For cell sorting, cells were harvested, washed three times with ice-cold PBS, pipetted through a Flowmi cell strainer (70 μM mesh size) and diluted to a final concentration of 200 cells/μL in ice-cold PBS. 400 nL of lysis buffer (100 mM triethylammonium bicarbonate (TEAB), 0.2% dodecyl β-D-maltoside (DDM), and 10 ng/μL trypsin/LysC) was dispensed into a 384-well plate (Eppendorf LoBind) using a Mantis liquid handling robot (Formulatrix). Single-cells were sorted and dispensed using the cellenONE platform (Scienion).

For protein digestion, the plate was incubated at 50°C for 1.5 hours with continuous rehydration using ultrapure water via the cellenONE system to prevent well evaporation. The plate was subsequently cooled to 10°C while maintaining rehydration. The digestion was quenched by addition of 5 μL of 1% trifluoroacetic acid (TFA) to each well.

For sample cleanup and loading, C18 Evotips (Evosep Biosystems) were prepared following the manufacturer’s protocol. Tips were conditioned by incubation in 1-propanol for 1 min, followed by two washes with 50 μL EvoB (0.1% formic acid (FA) in acetonitrile). Tips were then re-equilibrated with 1-propanol for 1 min and washed twice with 50 μL EvoA (0.1% FA in water). Washing steps were performed at 700 g for 1 min. Finally, 70 μL of EvoA was added as a pre-loading volume with a short spin (700 g, 10s).

Sample loading was performed using a Bravo liquid handling robot (Agilent Technologies), as previously described (27, 28). Briefly, samples were loaded onto prepared Evotips, washed with 10 μL of EvoA and also transferred onto the tips. Tips were subsequently spun at 700 g for 1 min, washed with 50 μL of EvoA, and stored with 200 μL EvoA maintained on top of the C18 material to prevent drying.

### Data-independent acquisition LC-MS analysis of single-cell experiment

Proteomics analysis was performed using an Orbitrap Astral mass spectrometer (Thermo Fisher Scientific) interfaced with an Evosep One chromatography system (Evosep Biosystems). Peptides were eluted from C18 Evotips using the Whisper Zoom 80SPD (16.3-minute separation) method and chromatographically resolved on an Aurora Elite column (5 cm × 75 μm i.d., 1.7 μm C18 particles, IonOpticks) maintained at 50°C in a custom heating module.

Mass spectrometric acquisition utilized a FAIMS Pro interface operated at −40 V compensation voltage with carrier gas flow set to 3.5 L/min and an EASY-Spray source (both Thermo Fisher Scientific). An electrospray voltage of 1,900 V was applied for ionization, and the radio frequency level was set to 40. MS1 survey scans were acquired from 380 to 980 m/z in the Orbitrap analyzer at a resolution of 240,000 (at m/z 200) with a normalized automated gain control (AGC) target of 500% and a maximum injection time of 100 ms. The Astral analyzer was operated in MS/MS data-independent acquisition (DIA) mode with a scan range of 150 to 2000 m/z covering the precursor range of 380 to 980 m/z with 45 variable-width isolation windows optimized on the basis of the precursor density generated via the py_diAID software^44^. The windows were acquired with 26 ms maximum injection time and an AGC target of 800%. The isolated ions were fragmented using higher-energy collisional dissociation (HCD) at 25% normalized collision energy.

### Single-Cell Raw MS data analysis

The single-cell raw files were searched together using alphaDIA version 1.10.3 in Python 3.11. Data was searched against a PeptDeep-predicted library from a mouse FASTA database (UP000000589, downloaded: 2024/01/17). For library prediction, a custom and optimized model by transfer learning was used for library prediction without fixed modifications, precursor length from 7 to 35 amino acids, precursor charge 2-4, precursor m/z 200-1800 and standard settings if not mentioned differently. A two-step search was used in AlphaDIA with the following parameters in the first search: target_ms1_tolerance = 4 ppm, target_ms2_tolerance = 7 ppm, target_candidates = 3, initial_ target_ms1_tolerance = 10 ppm, initial_ target_ms1_tolerance = 15 ppm and initial_rt_tolerance = 300, group_level was inferred on the basis of genes and the nn_hyperparameter_tuning was enabled; the second search parameters were the same with additional parameters: target_candidates = 5, batch_size for calibration set to 200, interference_strategy to ‘library’ and reuse_quant enabled to True. All precursors with run-level FDR of 1% and protein groups with global FDR of 1% were accepted. All other settings were used as standard (Supplementary Data 1).

### Mixed Species Raw MS data analysis

Raw mass spectrometry files from the previously published mixed species benchmark experiment^32^ were obtained from the ProteomeXchange Consortium repository (identifier: PXD046444). Data processing was performed using alphaDIA version 1.10.3 with the following parameters: library-free search was enabled, MS1 and MS2 mass tolerance were set to 5 ppm and 10 ppm respectively, and the precursor m/z range was restricted to 380-980 m/z. Protein identification was performed against a *H. sapiens*, *S. cerevisiae*, and *E. coli* FASTA file downloaded from UniProt in February 2024. For the directLFQ quantification, fragment ions were first filtered using a 0.9 XIC (extracted ion chromatogram) correlation cutoff with a minimum requirement of 12 fragments per protein. directLFQ was subsequently executed with the parameters: number of samples for quadratic fit (set to 50) and the minimum number of non-missing values (set to 3) (Supplementary Data 2). For the no filter dataset, we used the same settings as described above with the exception of the XIC set to 0 (Supplementary Data 3).

### Statistical Rationale QuantSelect algorithm

QuantSelect consists fundamentally of 3 modules: a preprocessing, deep learning and LFQ calculation module. The preprocessing step includes the feature extraction, feature engineering, preparation and standardization of fragment features acquired during alphaDIA search. During the feature extraction step the MS1 level intensity and ΔRetention time (ΔRT) and MS2 level intensity, mass error, XIC correlation and XIC height are retrieved from the alphaDIA framework. During the preprocessing step, new features are calculated including the mean correlation of intensity traces between each other, correlation of precursor-fragment traces and the number of missing values across fragments and samples. Next, all quality features are grouped by common protein groups represented by a three-dimensional tensor; dimension 1 represents the different fragment ions, dimension 2 represents the sample order and dimension 3 represents the quality feature kind (e.g. MS1 level intensity, MS2 level XIC correlation etc.). For training the 3-D tensor is flattened into a 2-D array where each row now represents a different fragment and each column represents a different feature. Intensity and height values are aligned during the preprocessing step using a re-implementation of directLFQ. The data is then standardized for each protein group separately. The deep learning module includes a neural network that consists of 4 fully connected layers, with batch norm 1D layers in between and a sigmoid activation function at the end. PyTorch v2.6.0 was used for deep learning. The algorithm was run with default settings using a regularization parameter (λ) of 0.8, trained for 40 epochs on 200 selected high-quality proteins. High quality proteins are defined by the largest number of identified fragments. The training process employed an 80:20 train-test split to prevent overfitting and ensure generalization to the full dataset. The model was then employed on the entire dataset and generated quality scores for each fragment. Fragments with a quality score ≥ 0.9 were retained in the dataset. The filtered fragments are aligned a final time and LFQ is calculated using directLFQ for the mixed-specie benchmark and single-cell data (Supplementary Data 4, 5).

### Validation of QuantSelect

To validate our loss function, we utilized a naive variance filter method where we calculate the standard deviation across fragments within a sample for each protein and excluded every fragment that exceeds the threshold of 3 times the standard deviation. We then calculated the fold changes and compared the naive variance filter method with QuantSelect’s implementation. The precision was here evaluated based on the standard deviation for each fold change within each species. The accuracy was evaluated based on the mean absolute error (MAE) with ground truth fold changes. In order to evaluate the quantification accuracy across the entire dynamic range, proteins were first binned in 1000 equally sized bins/abundance ranks. The MAE variance was calculated between each bin. A linear regression was then modeled on the abundance rank and to predict the MAE variance using the LinearRegression method from the sklearn v1.7.2 package. Additionally, the Pearson correlation coefficient and density of the data was calculated using the pearsonr and gaussian_kde method from the scipy v.1.16.2 package.

### Feature importance of deep neural network

Feature importance was calculated using a permutation-based method. Each input’s feature was permuted 30 times individually using the permutation method from numpy v2.1.0 package. After permutation, the input was fed into the model for prediction. The accuracy offset was used to determine the feature importance, greater offset to the un-permuted output equals greater importance. The offset was then normalized by division of the maximum offset.

### Comparison of QuantSelect and alphaDIA’s XIC_corr filter

Accuracy was evaluated based on the MAE of the intensity profile of each protein. Here, each protein profile was median normalized with ground truth profile for comparability. The MAE was then calculated as the mean absolute difference between ground truth and actual value.

In order to evaluate the average increase of offset between QuantSelect and XIC_corr filter, we utilized the linear regression method from statsmodels.api.OLS v.0.14.4 to model the MAE of QuantSelect to predict the MAE of the XIC_corr filter using the same method mentioned above. The slope of the model was then used to estimate the average offset increase.

To measure the reproducibility, we separated the dataset in each respective experimental group and calculated the coefficient of variation (CV) across each abundance rank. Abundance ranks are assigned by sorting protein based on their log2 intensity in 15 equally sized bins, where rising bin number represents increasing protein abundance. For each bin the CV was calculated.

### Statistical and downstream analysis

To evaluate the performance of differential expression detection in the mixed species experiment, we implemented an FDR calculation that exploited the known ground truth of the experimental design. P-values of known differentially expressed proteins (S. cerevisiae and *E. coli* proteins) were first sorted by their significance. For a given p-value cutoff *P_thresh_* (p-values that should be differentially regulated including *S. cerevisiae* and *E. coli* proteins) the FDR was calculated as follows:

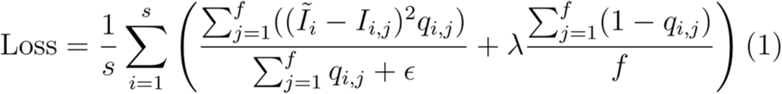

where *n_human_* (*P_thresh_*) represents the number of *H. sapiens* proteins with *p-values ≤ p_thresh_* and *n_total_ (p_thresh_)* represents the total number of proteins from all species with *p-values ≤ p_thresh_* The ROC AUC was calculated using the sklearn.metrics module from scikit-learn package version 1.7.1. For the unpaired two-tailed student’s *t*-test calculation we used the ttest_ind method from the scipy package v1.16.2. Multiple testing correction was done with the Benjamini Hochberg method using the multipletests method from the statsmodels v.0.14.4. The ROC curve and ROC AUC score were calculated based on the q-values using the roc_curve and roc_auc_score method from the sklearn v.1.7.2 package. Clusters were calculated using the hierarchical clustering method implemented in seaborn v.0.13.2 using the Euclidean distance as a distance metric.

## Results

### Regularized weighted variance as a loss function to solve the missing value problem

The aim of QuantSelect is to pick the optimal fragment ions in a quantitative DIA experiment, based on existing quality features using deep learning. Each fragment is characterized by a vector of quality-describing features such as ΔRetention time (RT), mass error, and extracted ion chromatogram (XIC) correlation that was calculated during processing with the search engine. This vector serves as input for a deep neural network, whose goal is to transform the n-dimensional input features into a single quality score that represents each fragment’s quantitative reliability, so as to classify fragments by their quality (Figure 1A). The output is a summarized quality score that can be transformed into a binary include/exclude decision.

**Figure 1.**
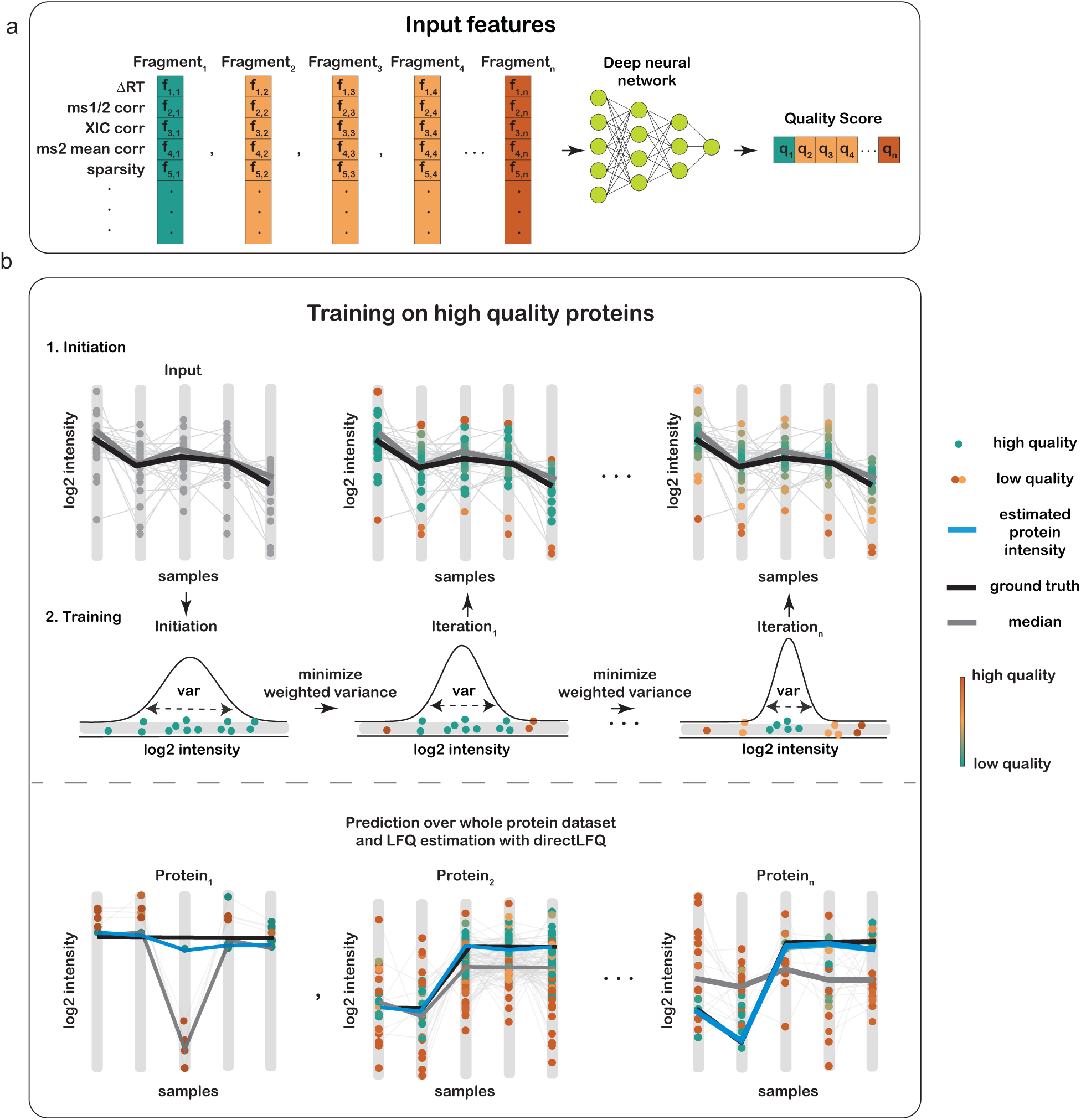
Overview of QuantSelect. **A)** QuantSelect uses a deep neural network to predict fragment quality scores from 11 features spanning MS1 and MS2 levels (see Methods). **B)** QuantSelect workflow. Training minimizes weighted variance of fragment intensities using high-quality proteins where the median trace approximates ground truth. The trained model then predicts quality scores across the full dataset to guide fragment selection.

A critical challenge in this approach is the absence of ground truth labels (29). QuantSelect addresses this by leveraging the variance as a proxy for quantitative quality under the assumption that fragment ion traces from highly abundant proteins exhibit similar intensity profiles on average (Figure 1B). However, this assumption requires fragment traces to be directly comparable, which we achieve through a preprocessing step that applies directLFQ normalization to correct systematic biases. For model optimization we utilized the weighted variance as a loss function as follows:

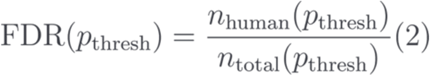

The optimization goal is to minimize the weighted variance across identified fragment ions within each sample for each protein. For each protein, we represent fragment intensities as a log2 matrix *I ∈ R^sxf^*, where *s* indexes samples and *f* fragments. The model assigns a quality score *q_s,f_* to each fragment, weighting its contribution to the estimated protein intensity *Ī_s_*, defined as the median fragment intensity within sample *s*. The loss function penalizes high weighted variance among fragments while applying a regularization term (controlled by λ) to discourage “lazy” convergence in which the network favors a single fragment and ignores the rest.

A small *ε* is added to the denominator to maintain numerical stability when weights are near zero. Training is restricted to high-quality proteins whose fragment intensities cluster around a consistent median; deviations from this median serve as a proxy for fragment reliability. Fragments with large quadratic deviations receive low scores, whereas those that closely follow the median are rewarded with high scores. This way the model iteratively minimizes the weighted variance of fragment intensities for each sample, progressively learning to recognize feature patterns that indicate fragment quality without seeing any intensity values. After training, the prediction is done on the entire dataset including low abundant proteins where the variance-based ground truth assumption may not apply.

### QuantSelect effectively scores fragment quality

We applied QuantSelect to a published mixed species dataset(24) consisting of 18 samples with predetermined ratios of *H. sapiens*, *S. cerevisiae* and *E. coli*, where fold changes are known for different sample comparisons (Supplementary Figure 1, Supplementary Data 6) consisting of comparison E5H50Y45:E45H50Y5 with a high fold change of 9 for both *E. coli* and *S. cerevisiae*, and comparison E10H50Y40:E40H50Y10 with a medium fold change of 4 for both *E. coli* and *S. cerevisiae* and comparison E20H50Y30:E30H50Y20 with a low fold change of 1.5 also for both *S. cerevisiae* and *E. coli*. The proportion of the *H. sapiens* sample remained constant in all comparisons.

Initial investigation of unfiltered fragment ions without QuantSelect revealed very low precision, characterized by large spread of fragments within each species, and low accuracy, reflected in the distance from true fold change (red dotted lines in Figure 2A, Supplementary Figure 2A, Supplementary Data 7). We then tested QuantSelect’s capabilities of classifying fragments by their quality. When calling QuantSelect with default settings, training with 200 proteins and 40 epochs completed in under 5 minutes, and revealed a bimodal distribution of quality scores (Supplementary Figure 3A, B, Supplementary Data 8,9). A cutoff at 0.9 resulted in 14.1 million fragments classified as low and 25.2 million fragments classified as high quality (Supplementary Data 9). The results showed that QuantSelect effectively separated fragments by quality, with low quality fragments having poor- and high-quality fragments having high precision and accuracy (Figure 2B, Supplementary Figure 2B, Supplementary Data 7). As mentioned earlier, we had used the quadratic deviation from the log2 median intensity (variance) to approximate the fragment’s quantitative quality of highly abundant proteins during training and generalize the model to low abundant and quantitatively challenging proteins (where the ground truth approximation is not applicable). To test whether the model generalized beyond this simple variance intensity approximation, we applied a naive variance filter by removing all fragments that crossed the threshold of median ± 3 standard deviations resulting in 13 million removed fragments (Supplementary Figure 4, Supplementary Data 10). Although a slight improvement in accuracy and precision over no filtering, this was clearly inferior to QuantSelect’s high quality fragments (Figure 2C, Supplementary Figure 2C, Supplementary Data 7). These results indicate that QuantSelect captures quality indicators beyond simple variance estimation and instead learns a more complex feature representation for fragment quality assessment.

**Figure 2.**
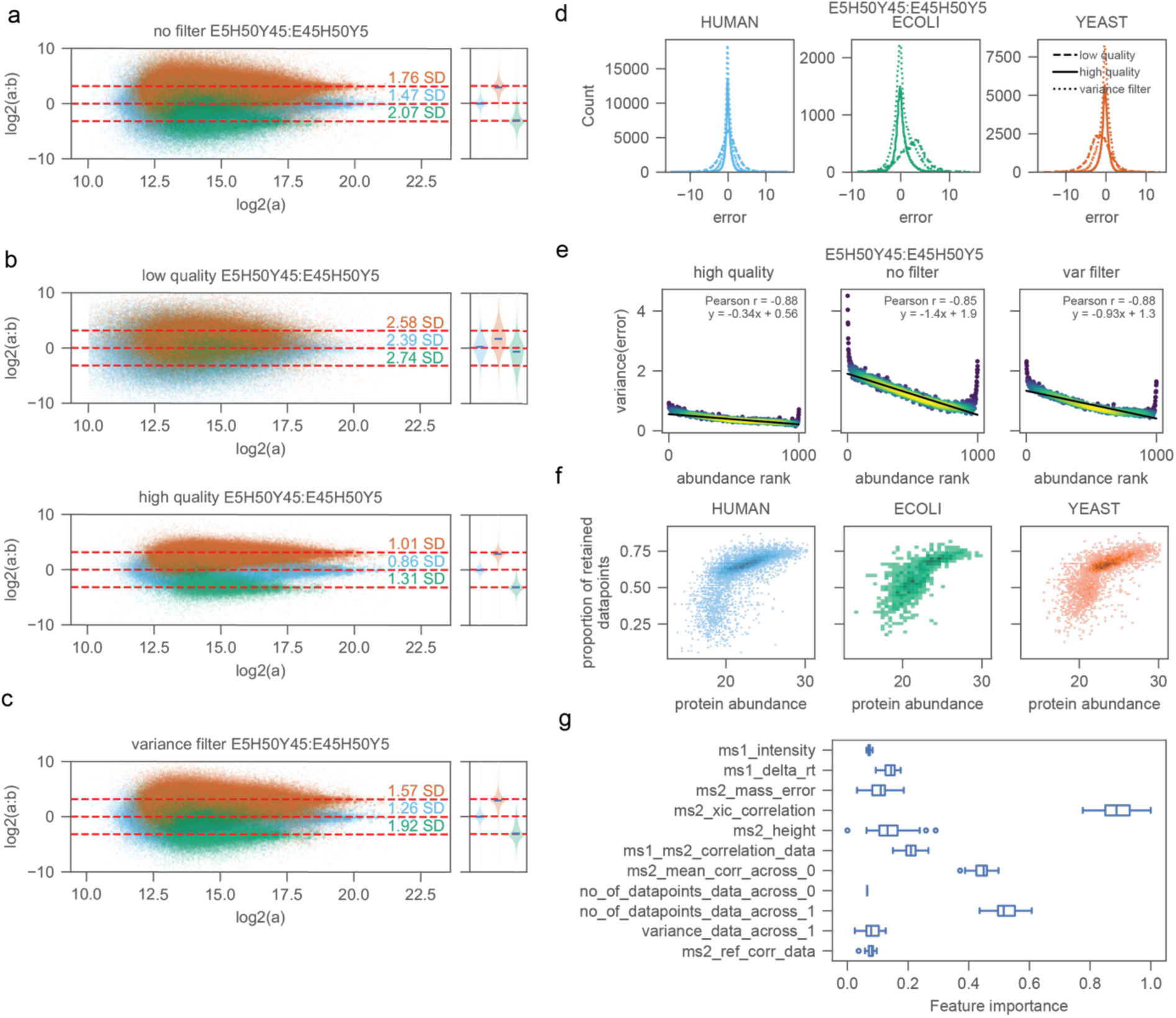
Investigating the efficacy of a deep neural network for fragment scoring. **A)** Distribution of individual fragments in a mixed-species benchmark (E5H50Y45:E45H50Y5) without any quality filtering. **B)** Same comparison after applying the QuantSelect quality score: upper, low-quality fragments; lower, high-quality fragments. **C**) Distribution after applying a naïve variance-based filter. Red dotted lines denote the ground-truth fold changes; standard deviations for each species are annotated. Blue represents *H. sapiens*, green represents *E. coli* and red represents *S. cerevisiae*. **D)** Accuracy of quantification shown as the distribution of deviations from the ground truth (error) for each species. **E)** Correlation between variance of absolute error and protein abundance rank. **F)** Relationship between protein abundance and the proportion of retained data points after QuantSelect filtering. **G)** Feature importance of QuantSelect’s neural network estimated by permutation analysis (n=30 permutations).

To assess these accuracies in more detail, we calculated the error for each log2 fold change and evaluated the precision of each method by the spread of the distribution (Figure 2D, Supplementary Figure 5A-B, Supplementary Data 11). QuantSelect’s high quality fragments were more tightly distributed around zero across all species, while low quality and variance-filtered fragments were more broadly distributed with greater deviations from the true fold change. The low-quality fragments even showed a systematic shift deviating from 0 error.

Low intensity fragments are often characterized by high levels of interference and thus lack accurate and precise quantification. To explore how QuantSelect alleviates this abundance-dependent quantification bias, we performed linear regression analysis between the variance of the error and abundance rank. This revealed that unfiltered fragments and naive variance filtered fragments had a marked linear relationship between abundance and quality score (slopes of 1.4 and 0.93 respectively), while high quality fragments showed less than half the slope (0.34), indicating a more stable quantification across the entire dynamic range as a result of a more discriminating fragment selection (Figure 2E, Supplementary Figure 6A-B, Supplementary Data 12).

QuantSelect’s quality scoring enables the assessment of the fraction of good versus bad fragments per protein. We plotted the fraction of fragments retained against protein intensity, which shows a weak linear relationship, which breaks down in the low abundance range (Figure 2F, Supplementary Data 13). Very low abundant proteins therefore have two challenges: fewer fragments and a larger fraction of bad fragments. However, even in the low abundance region, many proteins can still be quantified well by QuantSelect. This demonstrates that there is not a given intensity range where quantification is always poor, but that this strongly depends on the individual case. If multiple features are considered good quantification may still be possible.

Finally, we scored the importance of the individual features using permutation-based extraction. This showed that XIC correlation between individual fragment ions was most important, followed by mean correlation of all fragment traces (ms2_mean_corr_across_0) and number of datapoints. The remaining features were more balanced in their contributions (Figure 2G, Supplementary Data 14). The identification of XIC correlation as the most important feature aligns well with its importance in established DIA frameworks (6, 18, 23, 30). The fact that QuantSelect independently identified XIC correlation as the most important feature despite having no ground truth but relying solely on a self-supervised learning, demonstrates the effectiveness of our deep learning framework.

### QuantSelect improves precision and accuracy compared to single-metric XIC correlation filter

Next, we evaluated the performance of QuantSelect in combination with our directLFQ algorithm on the aforementioned published mixed species dataset. To this end we compared three fragment selection approaches: (i) no fragment selection (directLFQ-nofilter), (ii) QuantSelect-based fragment selection (directLFQ-QuantSelect), and (iii) the XIC correlation-based fragment selection implemented in the alphaDIA framework (directLFQ-XIC_corr), which applies an extracted ion chromatogram (XIC) correlation cutoff filter. For all three approaches, we used directLFQ for the subsequent LFQ calculation.

Across all three-fold-change comparisons (Figure 3A-C, Supplementary Data 15), QuantSelect consistently improved precision by 12-15% compared to both directLFQ-nofilter and directLFQ-XIC_corr, with the largest gains in challenging low-abundance proteins. Accuracy improvements were most pronounced at low fold changes, where QuantSelect achieved median values closer to ground truth across all species. Notably, directLFQ-XIC_corr sometimes performed worse than no filtering at all (Figure 3A-C), revealing a key limitation of single-metric cutoffs: for proteins with limited fragment coverage, stringent filtering excludes valid quantitative information that median-based aggregation would otherwise handle robustly.

**Figure 3.**
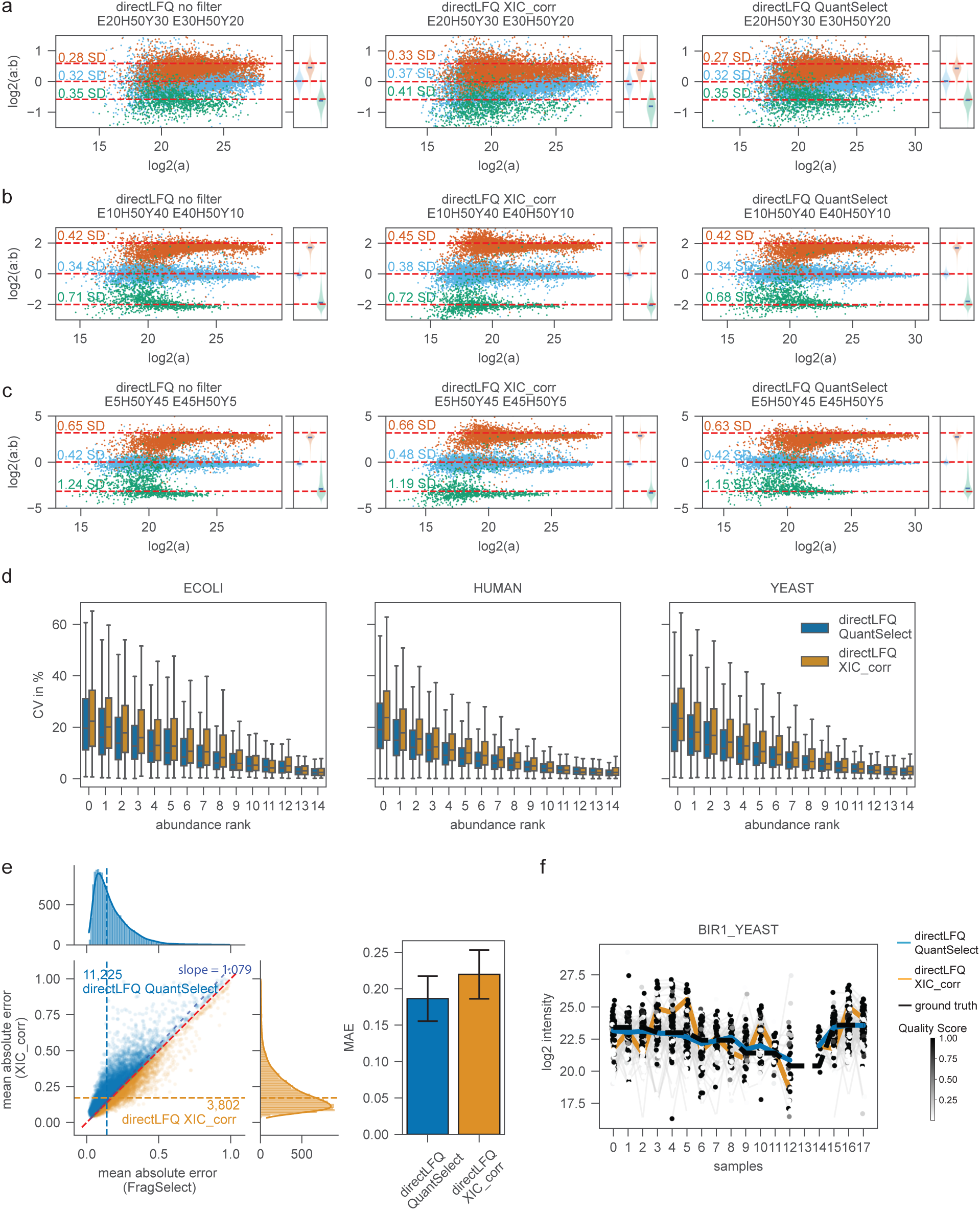
Comparing the quantitative performance of directLFQ-QuantSelect and directLFQ-XIC_corr on protein group level. Distribution of log2 fold change of fragments using directLFQ no filter (left), directLFQ-XIC_corr (middle) and directLFQ-QuantSelect (right) for the comparison E20H50Y30:E30H50Y20 **(A)**, E10H50Y40:E40H50Y10 **(B)** and E5H50Y45:E45H50Y5 **(C)**. The red dotted lines represent the ground truth. Standard deviations are annotated for each species distribution. Blue represents *H. sapiens*, green represents *E. coli* and red represents *S. cerevisiae.* **D)** Coefficient of Variation (CV) in percent of directLFQ-QuantSelect and directLFQ-XIC_corr across different abundance ranks. **E)** log2 deviation from ground truth for directLFQ-QuantSelect and directLFQ-XIC_corr. The horizontal dotted line (orange and blue) represents the median of either MAE (directLFQ-QuantSelect) or MAE (directLFQ-XIC_corr). The red line separates the plot in two parts; the upper part represents MAE(directLFQ-QuantSelect) < MAE(directLFQ-XIC_corr) and lower part represents MAE(directLFQ-XIC_corr) < MAE(directLFQ-QuantSelect). A linear regression was fitted against the data. The slope of the regression is annotated. The linear regression is plotted in blue dashes. Bar plot represents the mean MAE from ground truth of directLFQ-QuantSelect and directLFQ-XIC_corr. **F)** Log2 intensity trace of fragments against samples. Quality scores generated by directLFQ-QuantSelect are annotated according to the legend gradient. Black line represents the ground truth intensity trace, blue and orange represents directLFQ-QuantSelect’s and directLFQ-XIC_corr’s intensity trace.

Overall, alphaDIA’s filtering approach led to the exclusion of fragments with valid quantitative information, as evidenced by decreased precision in directLFQ-XIC_corr compared to directLFQ-nofilter. This effect was observed consistently across high, medium, and low fold change comparisons (Figure 3A-C), while QuantSelect demonstrated superior performance in balancing accuracy and precision. Specifically, QuantSelect showed marked improvements in precision across all fold change ranges (high, medium, and low) compared to both directLFQ-XIC_corr and directLFQ-nofilter and even improved accuracy for the most challenging low fold change comparison. These findings indicate that while single-metric cutoff approaches can improve quantification accuracy for a subset of proteins, they concomitantly deteriorate precision for others. QuantSelect addresses this limitation through a more intelligent fragment selection that maximizes the retention of informative fragments, thereby maintaining high precision while preserving adequate accuracy.

Having assessed precision and accuracy, we next evaluated quantitative reproducibility. When we binned proteins according to abundance, directLFQ-QuantSelect demonstrated better CVs than directLFQ-XIC_corr across all species, with particularly marked improvements by up to 15% in the low abundance range at the protein group level (Figure 3D, Supplementary Data 16).

To obtain a more comprehensive evaluation of quantification accuracy, we calculated the mean absolute error (MAE) as the absolute average deviation of individual protein group intensity traces from their ground truth values (Supplementary Figure 7, Supplementary Data 17). Linear regression analysis of the MAE of directLFQ-QuantSelect and directLFQ-XIC_corr yielded a slope of 1.079, which means that directLFQ-XIC_corr’s MAE has an increase of 7.9% on average compared to directLFQ-QuantSelect’s MAE (Figure 3E, Supplementary Data 18). We then bisected the scatter plot with a reference line separating regions where MAE (directLFQ-XIC_corr) < MAE (directLFQ-QuantSelect) (lower region) from those where MAE (directLFQ-QuantSelect) < MAE (directLFQ-XIC_corr) (upper region). The data points exhibited a greater density above the reference line, indicating that the numbers of proteins where the MAE (directLFQ-QuantSelect) < MAE (directLFQ-XIC_corr) with 11,225 proteins than for MAE (directLFQ-QuantSelect) > MAE (directLFQ-XIC_corr) with 3,802 proteins (Figure 3E).

The methodological advantages of QuantSelect are exemplified by representative proteins where the algorithm successfully identified and excluded low-quality data points that showed substantial deviation from ground truth values, despite displaying consistent log2 intensities across fragments. This selective fragment filtering was able to restore the true log2 intensities enabling accurate recovery of true protein quantification values compared to the XIC_corr method that missed the true values by orders of magnitude (Figure 3F, Supplementary Figure 8, Supplementary Data 19).

### QuantSelect increases sensitivity for differential expression analysis

To evaluate the performance of directLFQ-QuantSelect in differential expression analysis, we compared its sensitivity and accuracy against directLFQ-XIC_corr, across different fold change comparisons. At the protein group level, directLFQ-QuantSelect demonstrated higher sensitivity. For the high fold change comparison, directLFQ-QuantSelect identified 544 proteins at 1% and 3,687 proteins at 5% FDR compared to 329 at 1% and 2,197 at 5% FDR for directLFQ-XIC_corr, representing an up to 1.6-fold increase in sensitivity (Figure 4A, Supplementary Data 20, 21). This improvement was even more pronounced for the medium fold change comparison, where directLFQ-QuantSelect detected 1,180 proteins versus 295 for directLFQ-XIC_corr at 1% FDR and 3,693 and 2,933 at 5% FDR, up to 4-fold enhancement. For the low fold change comparison, both methods showed limited detection at 1% FDR (8 versus 11 proteins, respectively) due to the inherent challenges in identifying subtle expression differences. However, directLFQ-QuantSelect demonstrated substantially improved performance at 5% FDR (165 versus 11 proteins). We additionally evaluated the area under the receiver operating characteristic curve (ROC AUC) to assess the overall discriminatory power of each method independent of threshold selection. The ROC AUC analysis consistently showed improved AUC for directLFQ-QuantSelect over directLFQ-XIC_corr across all fold changes (Figure 4B, Supplementary Data 22). This is especially evident in challenging low fold change comparison where directLFQ-QuantSelect achieves a ROC AUC of 0.84 compared to directLFQ-XIC_corr ROC AUC of 0.72

**Figure 4.**
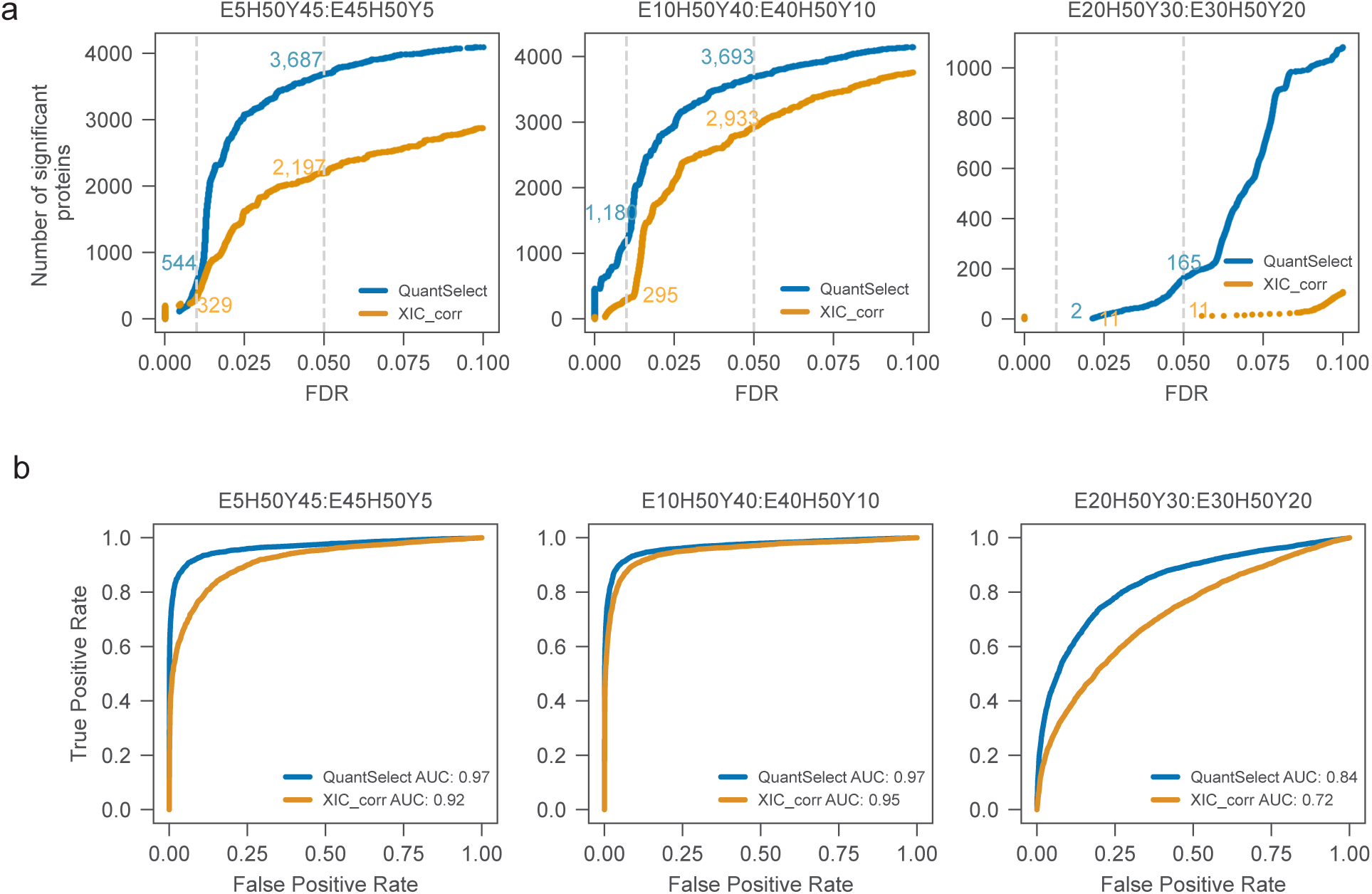
Comparing the sensitivity for differential expression analysis between directLFQ-QuantSelect and directLFQ. **A)** FDR against the number of significant proteins in directLFQ-QuantSelect and directLFQ-XIC_corr. Grey dotted lines represent 1 and 5% FDR. Annotation represents the number of significant proteins at 1 and 5% FDR threshold. N=3 replicates were used for two-tailed student’s *t-*test. **B)** ROC AUC curve using the q-value of directLFQ-QuantSelect and directLFQ-XIC_corr results.

### Applying QuantSelect to single-cell data

To evaluate both algorithms on a quantitatively challenging single-cell dataset, we analyzed embryonic stem cells from mice across three developmental stages: naive (ESC_n), formative (ESC_f), and primed (ESC_p). The dataset comprised 169 single-cells and a protein identification depth of 6,329 proteins. We ran QuantSelect with default settings using 40 epochs, an 80:20 train-test split. Our evaluation of coefficient of variation (CV) revealed that directLFQ-QuantSelect consistently decreased CV across the entire dynamic range compared to directLFQ-XIC_corr, when proteins were binned by log2 intensity in equal-sized bins (Figure 5A, Supplementary Data 23), demonstrating superior reproducibility. To further assess the reproducibility of biological groups, we calculated Pearson correlation coefficients between samples and performed hierarchical clustering analysis (Figure 5B, Supplementary Data 24). The clustering revealed consistent clustering of biological groups in directLFQ-QuantSelect, whereas the ESC_n population in directLFQ-XIC_corr segregated into two distinct clusters. directLFQ-QuantSelect identified more differentially regulated proteins (q-value<5%) in 2 out of 3 comparisons compared to directLFQ-XIC_corr (Figure 5C, Supplementary Data 25 and 26). This enhanced sensitivity was particularly evident when comparing naive and formative populations, where directLFQ-QuantSelect exclusively identified 378 significant proteins compared to 51 for directLFQ-XIC_corr, an overall increase of 18% in sensitivity.

**Figure 5.**
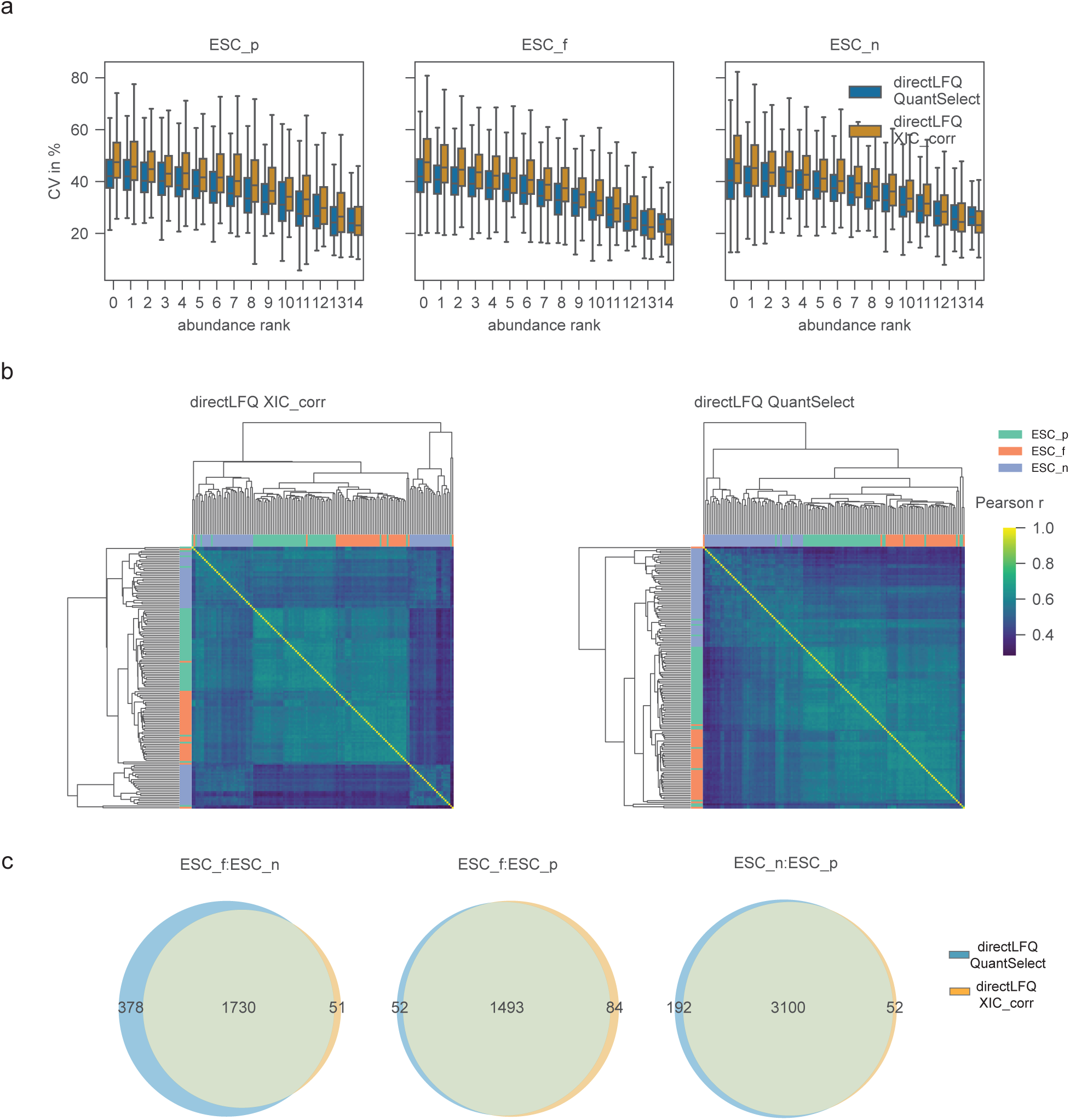
Comparing directLFQ-QuantSelect and directLFQ-XIC_corr on a single-cell dataset. **A)** Coefficient of Variation (CV) in percent of directLFQ-QuantSelect and directLFQ-XIC_corr across different abundance ranks. **B)** Heatmap of Pearson correlation data of directLFQ-QuantSelect and directLFQ-XIC_corr. Pearson correlation coefficient was calculated between each unique combination of samples. Hierarchical clustering with Euclidean distance was used for cluster analysis. **C)** Comparisons of significant proteins (q-value<5%) in comparison ESC_f/ESC_n (left), ESC_f/ESC_p (middle) and ESC_n/ESC_p (right) using directLFQ-QuantSelect and directLFQ-XIC_corr. A two-tailed student’s *t*-test were used for determining statistical significance with ESC_p n=57, ESC_f n=56 and ESC_n n=56.

## Discussion

The development of QuantSelect introduces a data-driven approach to fragment selection that moves beyond single-metric filtering. Traditional approaches have relied on cutoffs such as XIC correlation (6, 18). While computationally efficient, these methods fail to capture the complex factors that determine fragment reliability. Our results demonstrate that incorporating multiple features - including XIC correlation, retention time deviation, mass accuracy, and precursor intensities - provides a more robust foundation for identifying high-quality fragments.

The superiority of multi-feature assessment becomes particularly evident when examining proteins at the lower end of the abundance spectrum. Our analysis revealed that the linear relationship between retained fragments and protein abundance breaks down for low-abundant proteins, indicating that a single-metric e.g. intensity alone becomes an inadequate quality indicator. Similarly, alphaDIA’s XIC correlation cutoff improves accuracy for some proteins, but compromises precision for others. The single-metric filter fails to capture the nuanced quality indicators present in complex proteomics data. In contrast, QuantSelect’s neural network learns complex feature representation that enables more sophisticated quality assessment, resulting in consistent improvements in precision with adequate accuracy across the entire dynamic range.

A key innovation of QuantSelect lies in its versatility and adaptability. The framework’s modular design allows for straightforward incorporation of new features as they become available during data acquisition(16), making it inherently extensible. Any metric generated during the search process can be seamlessly integrated into the feature set, enabling continuous improvement as mass spectrometry technology evolves. Furthermore, the open-source nature of QuantSelect and its integration into the alphaDIA framework ensures accessibility to the broader proteomics community. The streamlined architecture of QuantSelect enables highly efficient computation. Training on our benchmark dataset, comprising millions of fragments and more than 15,000 protein groups required only 6 minutes from feature processing to intensity output generation. This computational efficiency makes QuantSelect accessible for standard desktop computers without specialized GPU hardware and can be readily used in conjunction with any downstream analysis tool (31–33).

The regularized weighted variance loss function introduced in QuantSelect addresses a fundamental challenge that extends beyond proteomics: the absence of ground truth in quantitative measurements. Traditional supervised learning approaches require labeled training data, which is often unavailable. Unsupervised clustering algorithms offer an alternative but become computationally intractable with increasing sample sizes and suffer from the “curse of dimensionality” as feature spaces expand (34). Our self-supervised approach circumvents these limitations by leveraging the inherent structure of the data - specifically, the principle that fragments from high-quality proteins should exhibit consistent intensity profiles(20).

Future developments could explore more sophisticated architectures. While our current implementation employs a simple fully connected neural network optimized for computational speed, alternative approaches such as transformer models (35) with additive attention mechanisms warrant investigation. Such architectures may yield performance improvements, though potentially at increased computational cost. Furthermore, QuantSelect uses the information provided by a search engine and does not itself calculate peak boundaries and peak picking(36). Optimizing this aspect of the pipeline will bring further benefits to QuantSelect and the overall quantification.

In conclusion, the principles underlying QuantSelect encompassing comprehensive feature utilization, self-supervised learning with a regularized optimization function, and adaptive quality assessment establish a new conceptual framework that can guide future developments in quantitative mass spectrometry, and potentially other analytical disciplines facing similar challenges.

## Data availability

The raw data can be downloaded via the ProteomeXchange with the following identifiers: mixed-species experiments PXD046444. The single cell experiment data will be released upon acceptance. All code to reproduce the results shown in this study is available under https://github.com/MannLabs/quantselect.

## Supplemental Data

This article contains supplemental data.

## Conflict of Interest

Matthias Mann is an indirect investor in Evosep Biosciences. The other authors declare no relevant competing interests.

## Acknowledgements

We thank our colleagues in the department of Proteomics and Signal Transduction for help and fruitful discussions. This study was supported by the Max Planck Society for Advancement of Science. M.O. was supported by the HORIZON-MSCA-2023-PF-01-01 project HETLEWY, no. 101151819.

## Author contributions

MT, EU, MO & MZ designed and performed the experiments; DTV, GW and CA developed the algorithm, DTV implemented the algorithm and the software package, DTV, MT, EU, CA, GW & MM analyzed and interpreted all data. CA & MM supervised and guided the project, DTV, CA, MM interpreted results and wrote the manuscript.

---

**Supplementary Figure 1:**
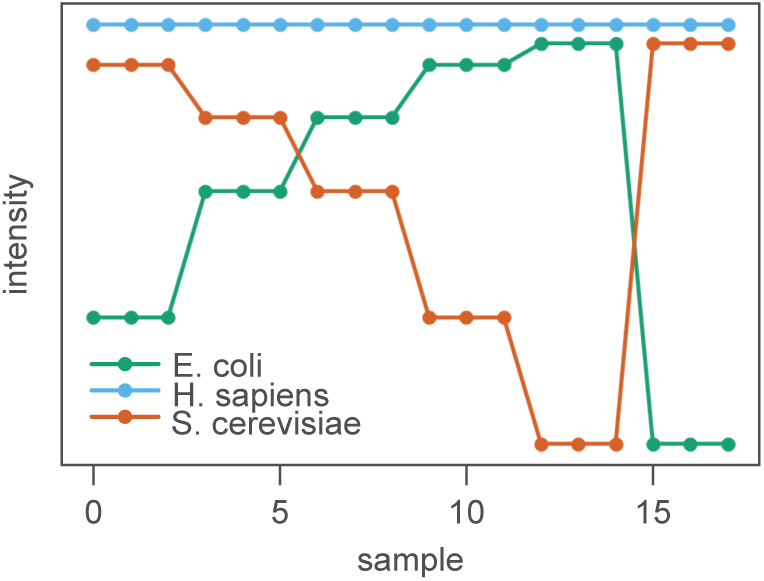
Illustration of intensity traces of mixed species experiment of *H. sapiens*, *S. cerevisiae* and *E. coli*. 200 ng of tryptic peptides were measured in six different ratios (E5H50Y45, E10H50Y40, E20H50Y30, E30H50Y20, E40H50Y10 and E45H50Y5). 3 consecutive measured samples are the equivalent of one technical replicate with the following ratios: 5:50:45, 10:50:40, 20:50:30, 30:50:20, 40:50:10 and 45:50:5 (*E. coli:H. sapiens:S. cerevisiae* ratio).

**Supplementary Figure 2:**
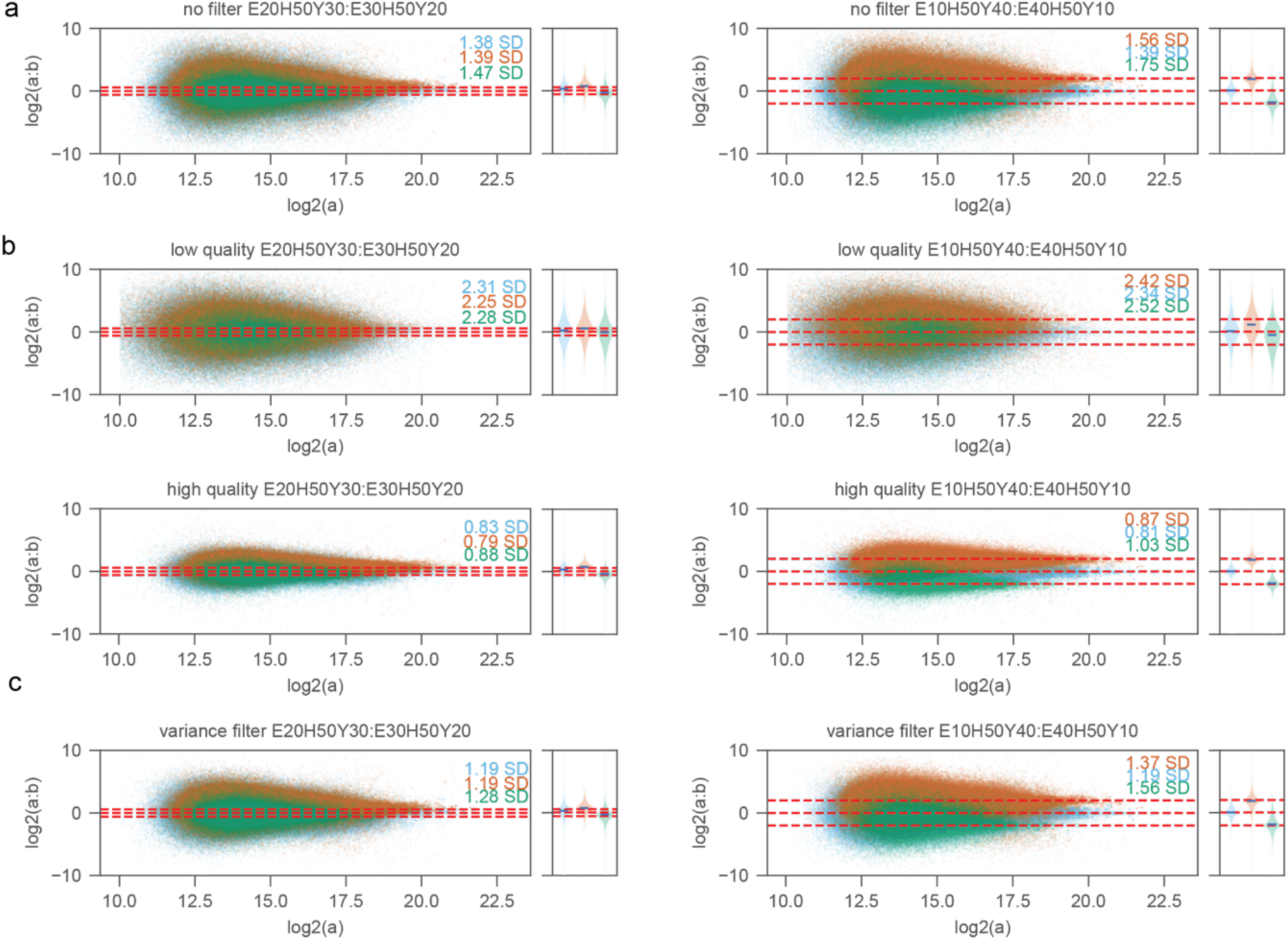
Distribution of log2 fold change of fragment ions for comparison E10H50Y40:E40H50Y10 (right) and E20H50Y30:E30:H50Y20 (left) of **(A)** no filter, **(B)** low quality (upper) and high quality (lower) and **(C)** variance filter. The left plot illustrates a scatter plot where each dot represents a fragment. The right plot illustrates violin plot. The red dotted lines represent the ground truth. Standard deviations are annotated for each species distribution.

**Supplementary Figure 3:**
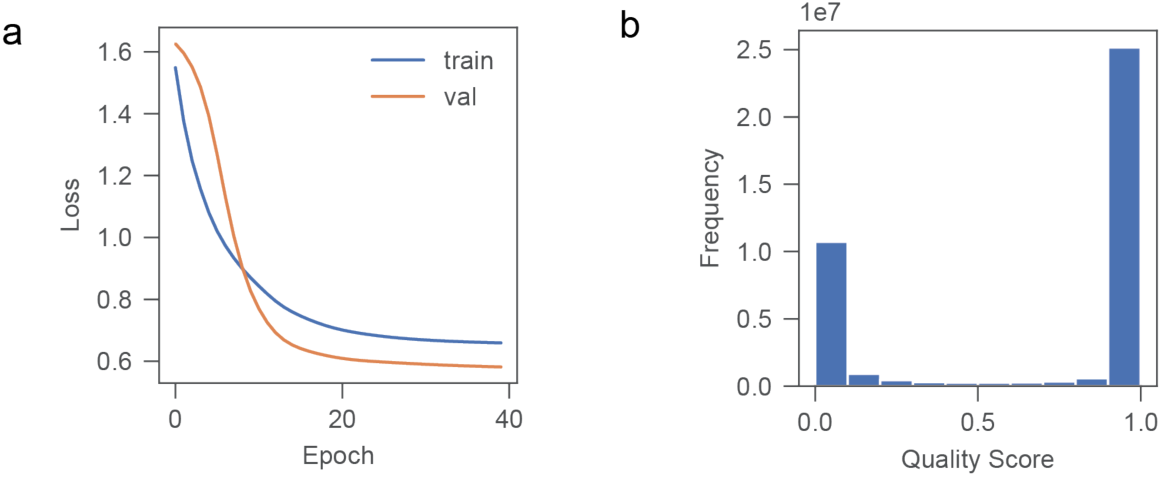
Training and prediction of QuantSelect’s model. **A**, QuantSelect was trained on 200 proteins for 40 epochs. Data was split into train and validation data using an 80:20 split. **B**, Distribution of quality scores generated by QuantSelect for each fragment.

**Supplementary Figure 4:**
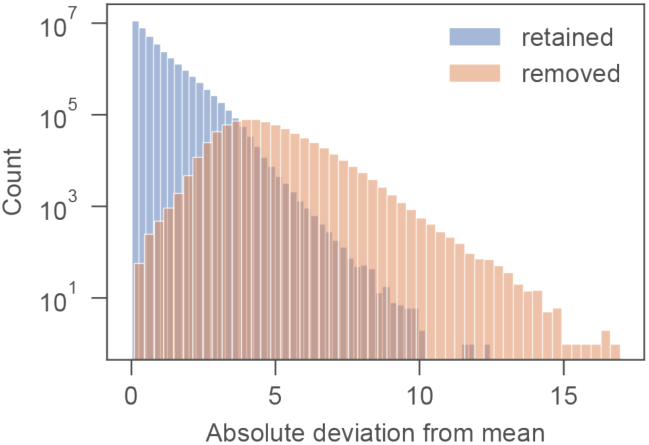
Distribution of absolute deviation from mean for each fragment during naive filter. Removed and retained fragments are colored in blue and orange respectively.

**Supplementary Figure 5:**
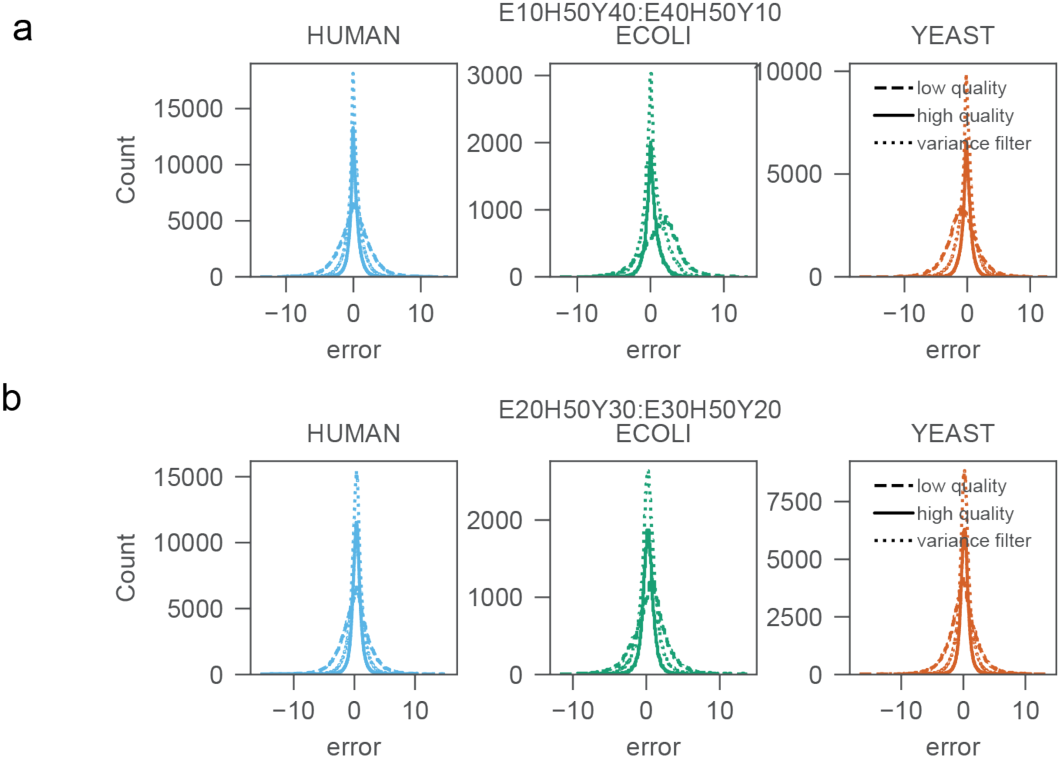
Distribution of MAE for each species for low and high quality and variance filtered fragments for comparison E10H50Y10:E40H50Y10 **(A)** and E20H50Y30:E30:H50Y20 **(B)**. Each linestyle represents either low quality, high quality and variance filtered fragments.

**Supplementary Figure 6:**
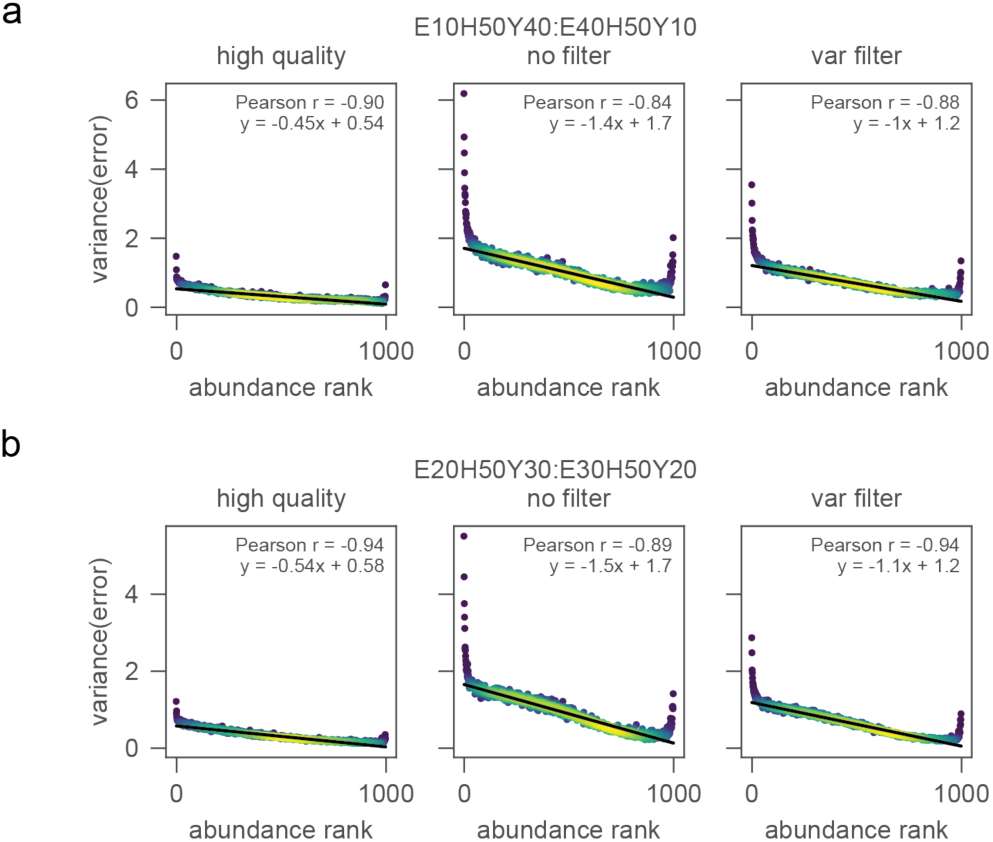
Correlation between variance of MAE for E10H50Y40:E40H50Y10 **(A)** or E20H50Y30:E30H50Y20 **(B)** and abundance. The abundance rank equals the bins calculated based on the log2 intensity with number of bins = 1000. A linear regression was fitted against the data. The Pearson correlation coefficient and linear regression equation are annotated. The density of the datapoints is illustrated by the gradient bar.

**Supplementary Figure 7:**
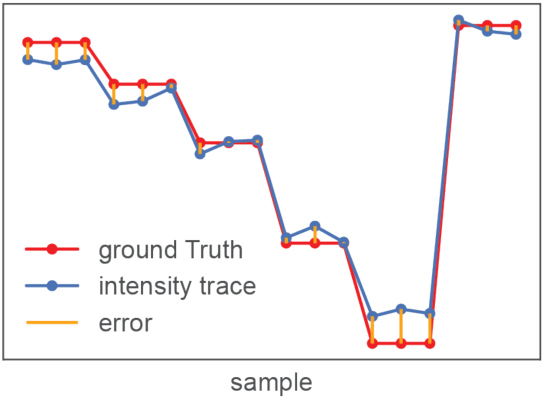
Illustration of mean absolute error calculation of intensity traces. Each trace of interest will be aligned to minimize the distance to the ground truth trace. The average of the absolute error will then be calculated and used as a measure of quantitative accuracy.

**Supplementary Figure 8:**
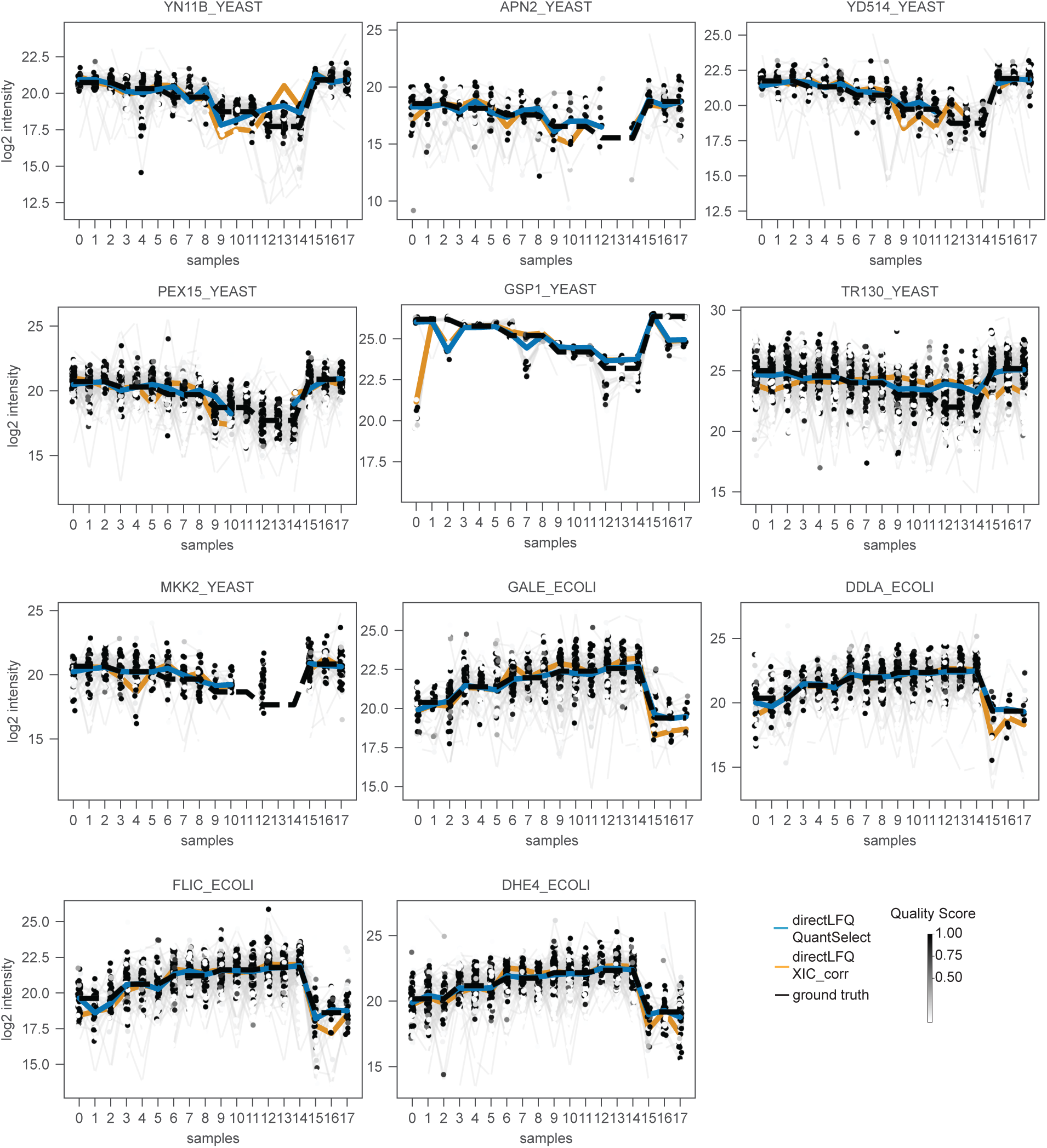
Example proteins where directLFQ-QuantSelect outperforms directLFQ-XIC_corr. Log2 intensity of fragments against samples. Quality scores generated by QuantSelect are annotated according to the legend gradient. The black line represents the ground truth intensity trace, blue and green represents directLFQ-QuantSelect’s and directLFQ-XIC_corr’s intensity trace respectively.

## Abbreviations

AGC: Automated Gain Control
AUC: Area Under the Curve
BSA: Bovine Serum Albumin
CV: Coefficient of Variation
DIA: Data-Independent Acquisition
ESC: Embryonic Stem Cell
FA: Formic Acid
FAIMS: Field Asymmetric Ion Mobility Spectrometry
FC: Fold Change
FDR: False Discovery Rate
GPU: Graphics Processing Unit
HCD: Higher-energy Collisional Dissociation
LC-MS: Liquid Chromatography-Mass Spectrometry
LFQ: Label-Free Quantification
MAE: Mean Absolute Error
mESCs: Mouse Embryonic Stem Cells
MS: Mass Spectrometry
MS1: First-stage Mass Spectrometry (survey scan)
MS2: Second-stage Mass Spectrometry (fragmentation)
m/z: Mass-to-charge Ratio
ppm: Parts Per Million
PSM: Peptide Spectrum Match
ROC: Receiver Operating Characteristic
RT: Retention Time
XIC: Extracted Ion Chromatogram

